# Phase transition pathways encode distinct physicochemical properties of biomolecular condensates

**DOI:** 10.1101/2025.03.26.645448

**Authors:** Xiaokang Ren, Leshan Yang, Michael Chen, Yifan Dai

## Abstract

The same intrinsically disordered proteins (IDPs) can form biomolecular condensates through distinct thermodynamic phase transition processes, such as temperature-dependent and osmosis-dependent pathways. These distinct thermodynamic driving forces should, in principle, induce phase transition by modulating different features of the solvent environments, so that different driving forces should correspond with distinct sequence grammars for phase transition. However, whether the molecular driving force is pathway-dependent and how these different pathways can define the properties and functions of condensates are largely unknown. Here, by employing a diverse set of solid-state and solution-state NMR techniques, we uncover that different phase transition pathways of the monomer unit of an IDP define the types of molecular interactions driving phase transition and stabilizing dense phases. By establishing a complete chemical shift profile of the IDP unit, we identified a unique interaction mode that specifically initiates the upper critical solution temperature transition process, an anion-dependent cation-pi interaction, in which Asp acts as a bistable molecular switch regulating a stepwise Arg-Tyr interaction in a critical temperature-dependent manner. We further show that the pathway-dependent molecular interactions encode condensates formed by the same IDP to exhibit different physical and electrochemical properties, which in turn enable distinct functions of condensates. Our study shows that besides the defined sequence grammar of a given IDP, the molecular driving forces under specific transition processes are different and can determine the structure and properties of condensates, which emphasizes an overlooked role of transition pathway on encoding the functions of condensates.

## Introduction

Phase transitions of biomacromolecules serve as the key thermodynamic mechanism mediating the formation of biomolecular condensates that regulate diverse cellular processes from transcription to stress responses^1–4^. Condensate formation has been found to regulate cellular functions through modulating the partitioning of biomolecules and solvent distribution. Partitioning of biomolecules can lead to specific cellular functions generated by sequestration or enrichment of specific sets of biomolecules, such as transcription enhancement on oncogenesis and protection of mRNA under heat or cold stress^5–17^. Modulating solvent conditions can lead to global cellular functions determined by the passive environmental effects created by solvent density transition^18–21^, such as altering cellular electrochemical equilibria, driving spontaneous chemical activities and buffering cellular water potentials^22–26^. These cellular processes are typically initiated by condensation driven by distinct phase transition pathways, specifically 1) temperature-dependent condensate formation in the case of heat or cold shock-dependent stress granule formations^10,14^ and 2) salt-dependent condensate formation in the case of hyperosmotic stress-induced condensates^27^.

Interestingly, though the thermodynamic driving forces underlying these transition pathways are different, the sequence grammars of the intrinsically disordered region of these proteins are highly similar as shown in the case of many highly repetitive and low-complexity IDPs (e.g, resilin-like polypeptides or FUS), which can both undergo salt-dependent and temperature-dependent phase transitions^28–30^. However, whether and how the transition pathway can determine the molecular driving forces mediating condensate formation and whether the pathway can determine condensate properties and functions are unclear (**Figure 1A**). Understanding this feature can shed light onto how distinct thermodynamic driving forces determined by the pathways can encode functions into condensates and potentially expand the molecular grammars of biomolecular phase transition in a pathway-dependent manner.

**Figure 1.**
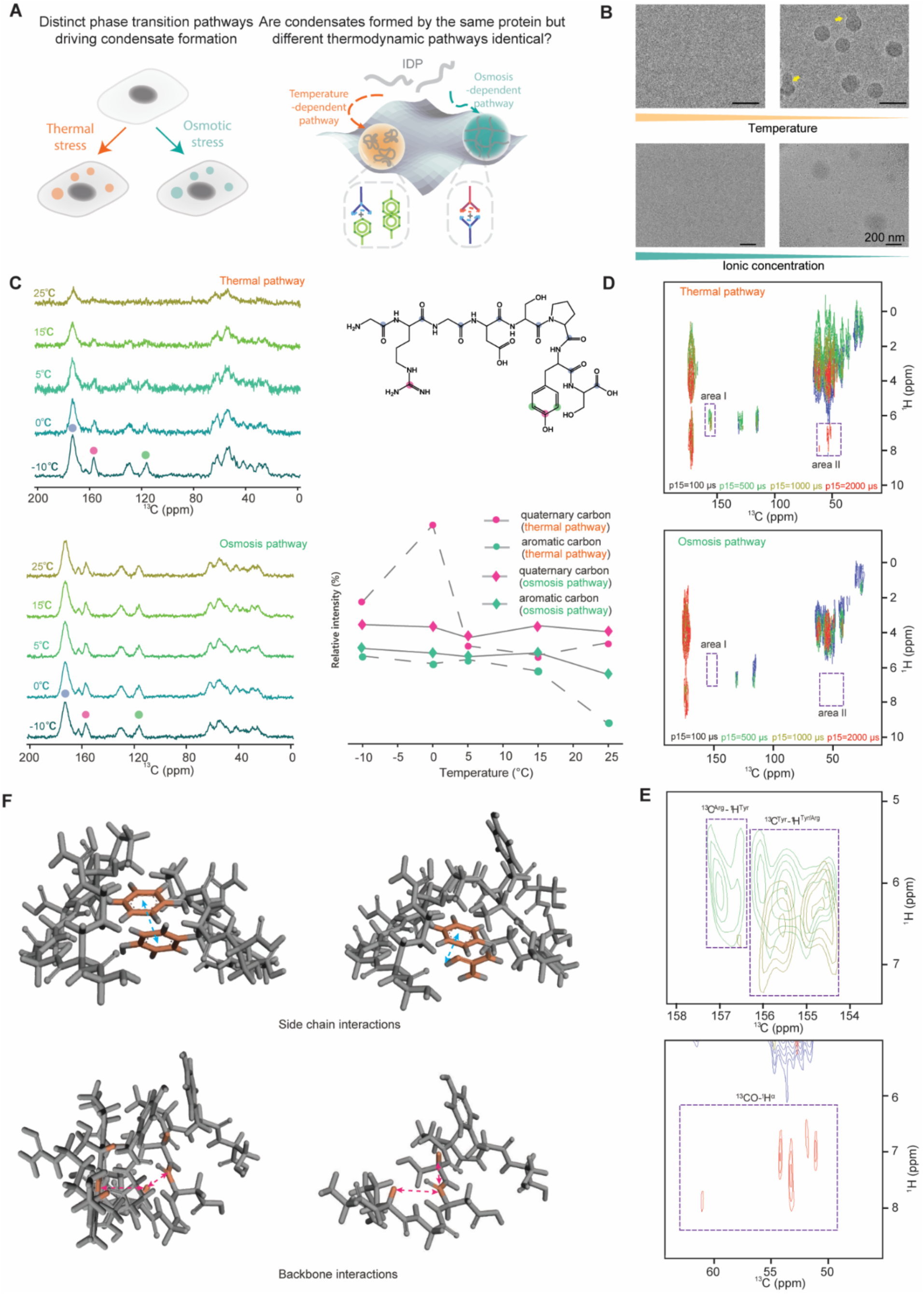
Solid-state NMR characterizations reveal the structural differences of condensates formed by distinct transition pathways. **A,** Biomolecular condensates inside cells can be driven by distinct environmental factors. The same IDP can undergo both osmosis-dependent (salt-dependent) and temperature-dependent transition processes. However, whether the condensates formed by the same sequence are the same is unknown. **B,** Representative Cryo-TEM images show the presence of condensates formed by distinct pathways. **C**, *In-situ* temperature-changing ^13^C CP-MAS was used to detect the conformational changes within different condensates from -10°C to 25°C. The red dots represent the carbon atoms directly attached to the phenolic hydroxyl group and the carbon atoms on the guanidine group, the green dots represent the carbon atoms in the ortho-position of the phenolic hydroxyl group, and the blue dots represent the carbon atoms on the peptide backbone and carboxylic acid group. And the relative intensity value of the selected carbon atoms NMR signal. The signal of the amide skeleton carbon atom is used as a standard. The meaning of each color is the same as in the picture above. **D,** ^1^H-^13^C HETCOR spectra of condensates formed by cooling or diluting with different contact times (100 µs, 500 µs, 1000 µs, and 2000 µs). **E,** Detailed structural information of the residue groups and the backbone based on merged cross points of cooling-inducing condensates. The marked area represents the interactions between tyrosine to arginine groups, tyrosine groups to tyrosine groups (Area I), and carbonyl carbon on the peptide backbone with hydrogen on amino acids α carbons (Area II). **F,** Scheme of the side chain interactions and backbone interactions that existed within the temperature-dependent condensates.

We herein first evaluate the molecular driving forces under 1) temperature-dependent and 2) salt-dependent phase transitions of the monomer unit of resilin-like polypeptide (RLP) (Gly-Arg-Gly-Asp-Ser-Pro-Tyr-Ser) using solid-state and solution-state NMR characterizations. We found that the molecular interactions through temperature-dependent and salt-dependent phase transitions are different, which demonstrate that transition pathways can determine the internal structure and molecular conformations within the condensates. With diverse solution-state NMR techniques, we established the complete chemical shift profile of the peptide at an atomic level. We then discovered that for temperature-dependent transition pathway, a stepwise phase transition process is mediated by a specific intra-chain interaction, in which an anion-dependent cation-pi interaction serves as a bistable switch. This switch occurs at a specific temperature, which coincides with the critical temperature of phase transition. We further verified this process in the full length RLP protein. Lastly, we show that the transition pathway can encode the physical and electrochemical properties of condensates by demonstrating that the internal dynamics of full length RLPs and the electrochemical environments vary between condensates formed through different pathways. These features further enable the condensates to exhibit pathway-dependent inherent chemical activities. Our study demonstrates that different thermodynamic transition pathways of the same IDP can lead to distinct molecular interactions, resulting in condensates with different properties and functions despite being assembled by the same biomolecular component. This finding demonstrates that the transition pathways can serve as an orthogonal factor, alongside sequence grammar, to control the biological functions of condensates.

## Results

To study distinct phase transition pathways using the same protein, we implemented the monomer of the widely studied RLPs (**Figure S1**), GRGDSPYS, which possesses all the key phase transition driving forces (i.e., cation-pi, pi-pi, electrostatic) as discovered in previous works^28,31,32^. This sequence can undergo both salt-dependent and temperature-dependent (i.e., UCST phase behavior) condensate formation, thereby allowing us to study the molecular driving force of distinct phase transition pathways using the same sequence. The short peptide unit is sufficient to drive phase transition, and notably, compared to complex long disordered protein, this peptide unit offers an incomparable opportunity to attribute chemical shifts into the atomic level of each residue, thus allowing us to construct a complete interaction profile to study the transition process.

### Transition pathways define distinct dense phase interactions

We implemented two distinct phase transition strategies to prepare condensates formed by the RLP monomer by 1) decreasing the ionic strength of the solution by dilution and 2) decreasing the temperature of the solution. We verified the formation of condensates based on these pathways using cyro-TEM (**Figure 1B**). We next separated the dense phases through centrifugation and flash froze the dense phases for solid-state NMR characterizations^33–38^. This allows us to directly evaluate the differences in the dense phase interactions within the condensates.

We first set out to study the dynamics of dense phase interactions using the *in-situ* temperature-changing ssNMR measurements by monitoring chemical shifts within the dense phases formed by different pathways under an external temperature gradient during NMR measurements. The changes in chemical shift during temperature change represents the motility of the peptide unit (**Figure 1C**). The resonances around 60-40 ppm were attributed to the aliphatic carbon atoms, the resonances around 160-100 ppm were attributed to the aromatic carbon atoms and unsaturated carbon atoms and the resonances around 180 ppm were attributed to the carbon atoms of the amide bond on the peptide backbone and carboxylic groups^39–41^. We found that at -10 °C, the chemical shifts within the dense phases formed through different pathways are identical, confirming that the condensates are formed through intermolecular interactions of the same monomer unit. However, as the testing temperature increases, the signal intensity from the dense phase formed by the cooling process continues to decrease. In contrast, the signal intensity of the dense phase formed by decreasing ionic strength remains unchanged (**Figure 1C**). Since ssNMR uses a cross-polarization/magic angle spinning (CP/MAS) method to detect signals, a rigid stacking pattern reflects on the improved efficiency of NMR acquisition compared to a flexible stacking mode ^42,43^. Therefore, as the testing temperature increases, the weakening of the resonances from the dense phase generated through the cooling pathway suggests that the molecules within the dense phase are flexible and the arrangement of molecules is transformed to a more heterogeneous disordered state^44^. This implies that the internal structures of the condensates formed by distinct pathways are different. We next selected the meta-carbon atoms on the tyrosine phenolic hydroxyl group (green dot), the carbon atom directly linked to the tyrosine hydroxyl group (red dot), and the quaternary carbon atom on the side chain of arginine (red dot) as the representatives of different side chain groups. The analysis of the strength of ssNMR signal to varying temperatures after normalization showed that although the increase of temperature drove the motion of the peptide into a more dynamic mode, the responses of different side chain groups in the dense phase to the temperature change were not the same. Specifically, the resonance intensity decreased to the 14.37 % of Arg residue and 0% of Tyr residues when temperature increased from -10 °C to 25 °C.

To illustrate the details of the dense phase interactions, we next implemented the 2D ^1^H-^13^C heteronuclear chemical shift correlation experiments (HETCOR) under magic angle spinning (MAS) to elucidate the intermolecular interactions within these dense phases formed by distinct pathways. Different contact times (p15) were used to ensure the detected distance from near (about 1.0 Å) to far (about 5.0 Å)^45,46^. In contrast to the condensates obtained by decreasing ionic strength (**Figure 1D**), distinct hydrogen to carbon-related regions were found in the cooling-induced condensed phase (**Figure 1E**). One region is due to the interaction between residues of tyrosine to arginine and tyrosine to tyrosine, and the other region is due to the interaction of carbonyl carbon on the peptide backbone with hydrogen on amino acids α carbons (**Figure 1E&F**). Besides, by analyzing regions of pi-based interactions, we found a split of the proton signal resonance only in the cooling-induced condensates (**Figure 2A**), implying the ring-current effect from aromatic rings **(Figure 2B**)^47,48^. These observations confirm that temperature-induced condensate formation triggers the conformational changes of the sidechain and backbone driven by cation-π and π-π interactions, which in turn stabilizes the dense phase. Taken together, the results from solid-state NMR provide direct evidence showing that the transition pathway can encode distinct molecular interactions in the dense phase.

**Figure 2.**
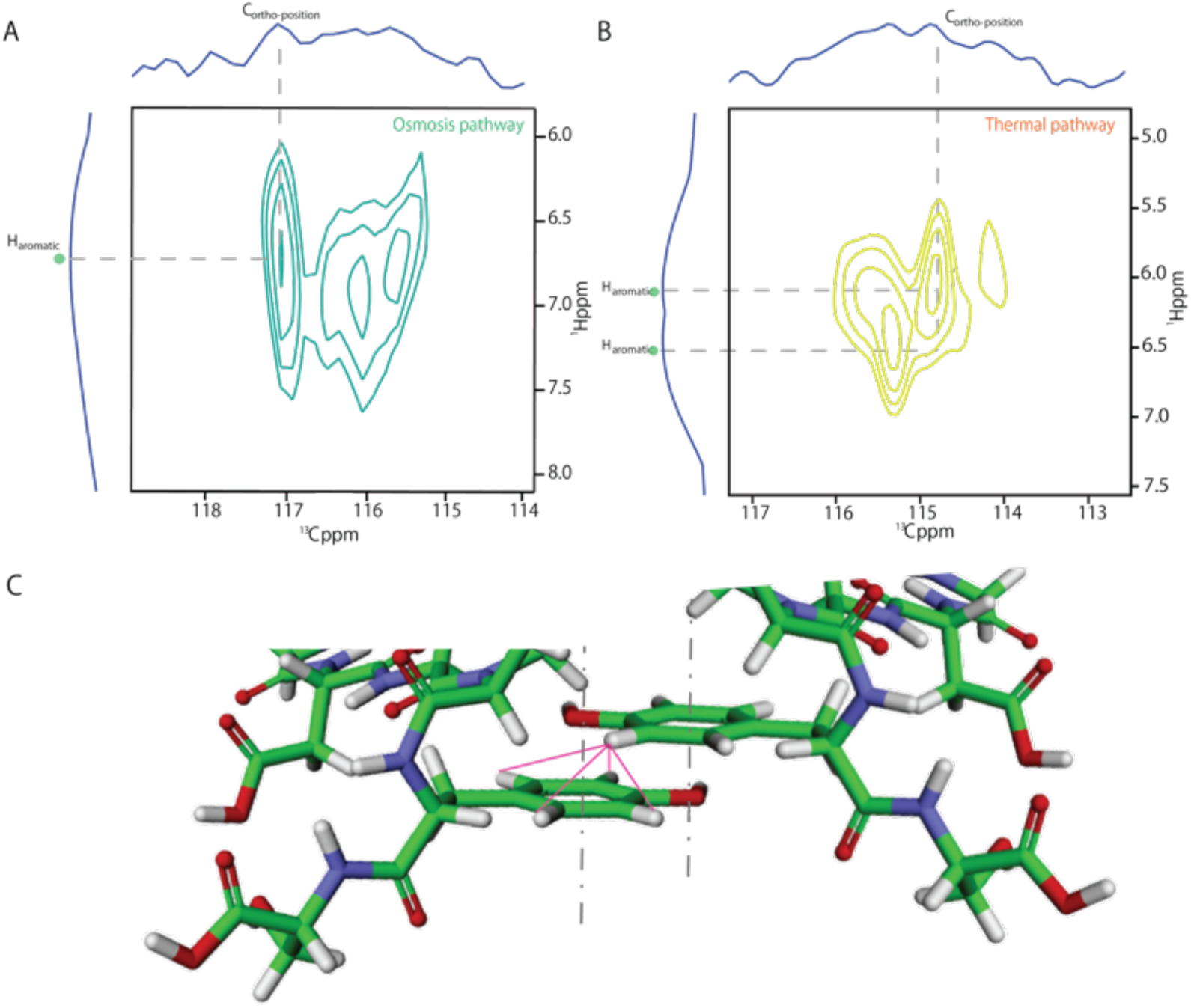
2D contour plot of the ^13^C-^1^H HETCOR correlation experiment for condensates formed by distinct pathways. **A,** 2D contour plot for condensates formed through salt-dependent pathway. The horizontal and vertical projections show the corresponding 13C and 1H spectra, respectively. **B,** 2D contour plot for condensates formed through temperature-dependent pathway. The horizontal and vertical projections show the corresponding 13C and 1H spectra, respectively. Dashed lines indicate the spatial connectivity along with the aromatic hydrogen atom and the ortho-position carbon atoms in phenol residues. In contrast to figure A, two aromatic resonances are clearly identified, which were induced by the ring-current effect caused by the aromatic π-π packing module. The **C**, Schematic structure of the above model. The pink lines represent the ring currents associated with aromatic moieties.

### Temperature transition is driven by a specific interaction based on anion solvation-dependent cation-pi interaction

To enable detailed understandings of the temperature-dependent transition process, we first used a series of solution-state NMR characterizations (**Figure S2**)^49,50^ to establish an atomic-level resolution profile of the chemical shifts within the RLP monomer (**Figure 3**). We next used the 1D *in-situ* temperature-changing ^13^C NMR spectra to determine the conformational changes of different amino acids in the temperature-induced condensate formation process. We found that distinct segmental dynamics were observed within the peptide (**Figure 4A**). As the temperature decreases from 37 °C to 4 °C, the chemical shift of the carbonyl carbon atoms in the peptide backbone shifts. Specifically, the chemical shift of the carbon in the Arg residue changes by 0.26 ppm, in the Asp residue by 0.18 ppm, in the Ser residue by 0.09 ppm, and in other residues by 0.01 ppm. Thus, the peptide can be divided into two segments: Gly-Arg-Gly-Asp, which possesses better conformational flexibility, serving as the promoter of the coil-to-globule transition process. Meanwhile, the Pro-Tyr-Ser segment experiences greater steric hindrance, resulting in slower translational movement during the transition process ^51–53^. These observations suggest that distinct segments within the peptide contribute to the UCST phase behavior of the RLP monomer differently.

**Figure 3.**
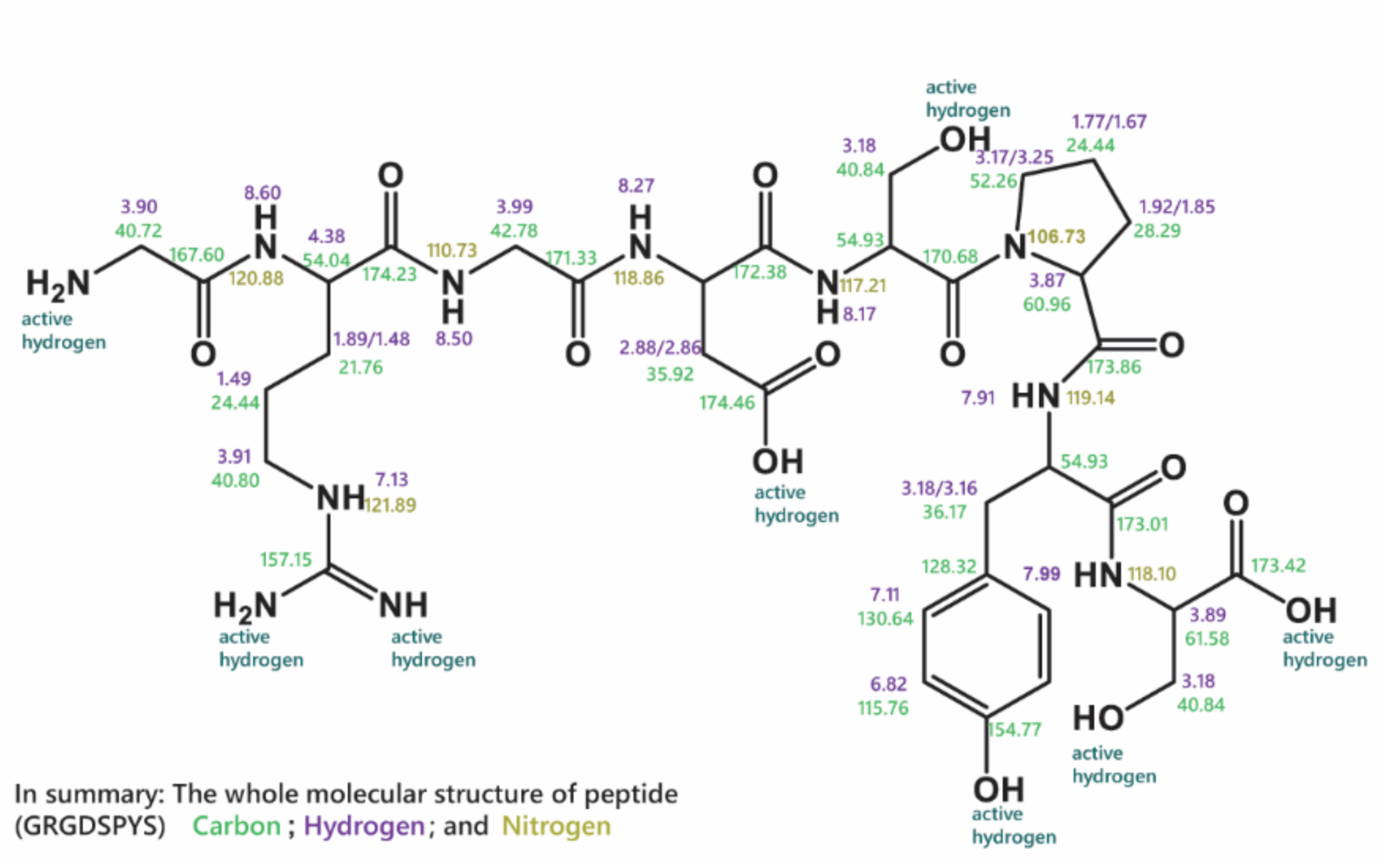
Complete chemical shift assignment of the monomer unit. Structural information based on the atomic level profile of the chemical shifts of the RLP monomer unit (GRGDSPYS) at 37 °C established by multiple solution-state NMR experiments.

**Figure 4.**
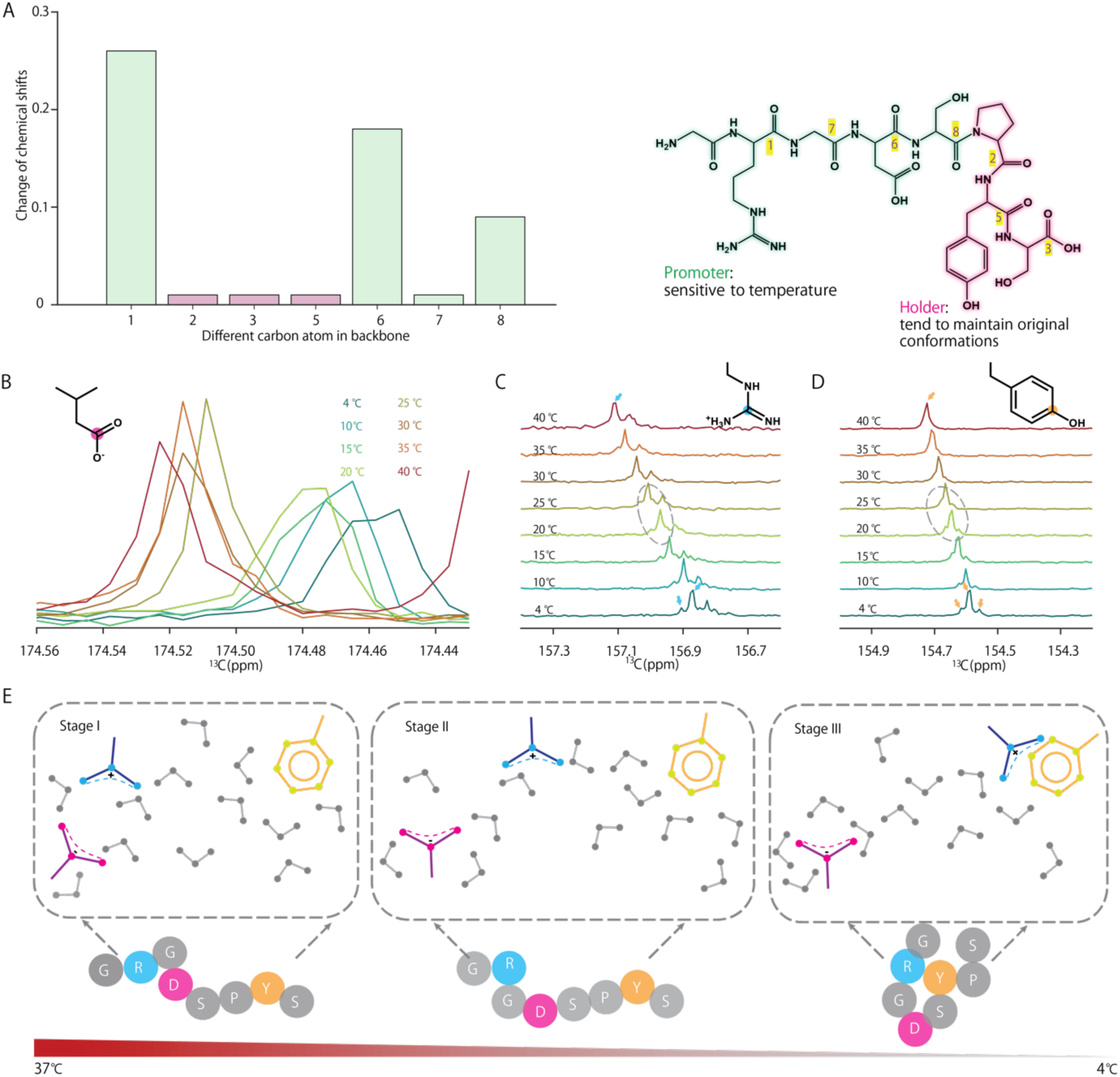
The solution-state NMR characterization reveals the underlying mechanism of the temperature induced formation of condensates through UCST transition. **A,** In-situ temperature-changing ^13^C NMR was used to detect the conformational changes of RLP monomer from 4 °C to 40 °C. The change of chemical shifts are represented by bar graphs for different carbon atoms on the backbones. Two distinct regions within the peptide were identified as a promoter for phase transition and a holder that maintains the conformational rigid of the interaction residues. **B,** The chemical shift of the carbon atoms within carboxylic groups at different temperatures. **C,** The chemical shift of the carbon atoms of guanidine groups at different temperatures. **D,** The chemical shift of the carbon atoms of the phenol groups of tyrosine at different temperatures. **E,** A proposed mechanism of the temperature induced condensate formation driven by anion-dependent cation-pi interactions.

To understand the specific driving forces, we next evaluated the contributions of electrostatic and pi-based interactions by analyzing the chemical shifts of positively charged Arg residue, negatively charged Asp residue, and aromatic-based Tyr residue during temperature transition ^54–56^, which serve as the critical “sticker” residues driving phase transition^31^. We found that the chemical shifts of Y and R residues show a monotonic shift to the low-frequency region upon decreasing temperature (**Figure 4B, C, & D**). The chemical shift for Tyr (orange dot) changed from 154.72 ppm to 154.58 ppm, while the chemical shift for Arg (blue dot) shifted from 157.11 ppm to 156.83 ppm. However, the chemical shift of Asp group (red dot) exhibited a bistable state, shifting from 174.51 ppm to 174.45 ppm during temperature transition (**Figure 4B**). This behavior indicates a critical point of conformational transition of the Asp residue between 20°C and 25°C^57^. Notably, the resonances of both Tyr and Arg split at the same temperature range, suggesting that a new conformational structure began to form in the surrounding environment^58,59^. This can be explained by the temperature-dependent free energy of hydration of the Asp residue^60^, which leads to the chain collapse driven by the competing interactions between Asp, water molecules and Arg. This interesting observation uncovered that the bistable Asp conformations serve as a molecular switch to initiate the cation-π interaction between Tyr and Arg and the temperature-dependent transition process can be initiated by a temperature-dependent solvation property of a specific residue (**Figure 4E**). These results suggest that UCST phase transition is a stepwise process involving enthalpic contributions from solvation, intermolecular interaction and water reorganization^61^.

### Pathway defines the diffusion dynamics of the peptide and full-length RLP_WT_ in condensates

As the interior interactions of condensates formed by distinct pathways are different, we next wondered whether these differences in interactions could encode condensates with distinct physical and chemical properties (**Figure 5A**). For the physical property, we reasoned that as the distinct interactions in the dense phase define the conformation of the chain, which defines the radius of the gyration of the chain^62^, the diffusion dynamics of the chain might be different. For the chemical properties, we reasoned that as the distinct interactions in the dense phase enforce different residues in the dense phase to define the asymmetry of ion and water partitioning^25,61,63,64^, the microenvironment of the condensates can be different, thereby enabling distinct capabilities to drive their inherent electrochemical functions^24,65^.

**Figure 5.**
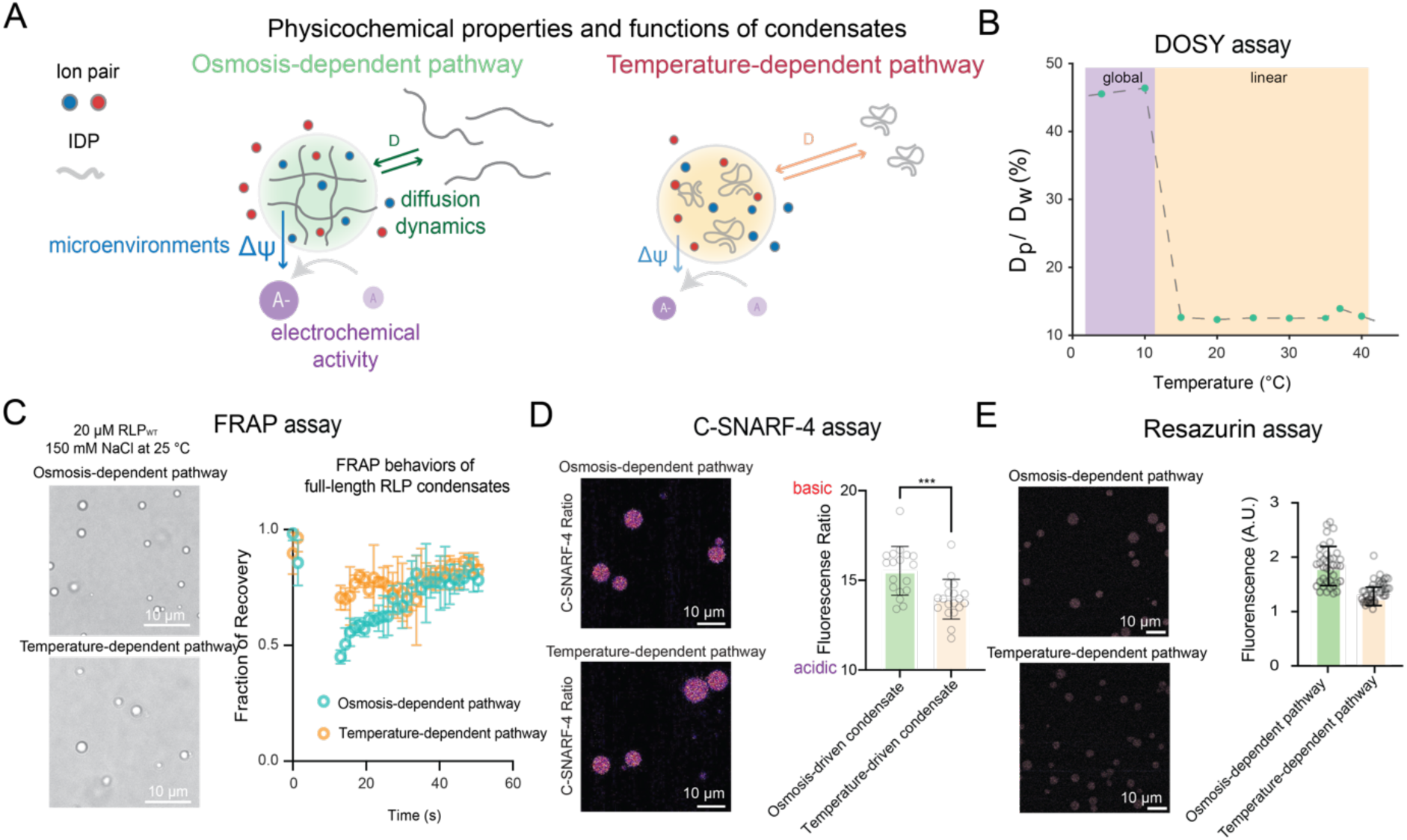
Pathway-dependent physicochemical properties and electrochemical functions of biomolecular condensates. **A,** Schematic illustration of how distinct reaction pathways influence the interfacial potential and diffusion coefficient within biomolecular condensates. **B,** In-situ temperature-changing 1H DOSY was used to characterize the molecular diffusion dynamics of the peptide in reference to water molecules. **C,** FRAP analysis of condensates formed by full length RLP_WT_ through distinct phase transition pathways. The final environmental states of the condensates formed by distinct pathways are identical. **D,** C-SNARF-4 assay measurement of apparent pH in the dense phase of condensates formed through different pathways. Higher the fluorescence ratio suggests a more basic pH. Each data point represents an individual condensate. Scale bar: 10 μm. **E,** Resazurin assay for the evaluation of the electrochemical activity of condensates formed through different pathways. Resazurin was incubated for 30 min prior to confocal imaging. Each data point represents an individual condensate. Scale bar: 10 μm.

We first investigated whether distinct pathways could encode different physical properties of condensates. To this end, we used the *in-situ* temperature-changing NMR diffusion spectroscopy (DOSY) to evaluate the diffusive property of the monomer RLP. We compared the relative diffusion coefficient of the peptide by referencing the proton signal of the aromatic group on the peptide to the proton signal of water molecules, through which we were able to identify the temperature (10-15 °C) at which condensate forms (**Figure 5B**). This temperature is different from the critical point temperature (20-25 °C) at which Asp undergoes conformational transition within the peptide. Interestingly, we found that the diffusion coefficient of the peptide increases upon decreasing the temperature, suggesting a coil-to-globule transition (**Figure S3A&B**). From ^1^H-^1^H NOE spectrum **(Figure S3C**), we also observed temperature-dependent cation-pi interaction between Arg and Tyr, which mediates the coil-to-globule transition ^66,67^. These observations further confirm that UCST phase transition is a multi-step process that delivers a compact peptide conformation, which mediates a higher free volume within the condensates, thereby contributing to enhanced diffusion after phase transition.

We next expressed the full-length RLP_WT_ ([GRGDSPYS]_20_)^19,29^ and evaluated whether such temperature-dependent multi-step process is also valid when the monomer is polymerized into a repeat of homopolymer. The final solution condition of the condensates prepared by distinct pathways is the same (see Methods for details). Using the *in-situ* temperature changing DOSY experiments, we found that the full-length RLP_WT_ also undergoes a similar multi-step transition process (**Figure S4A).** A noticeable proton signal of Arg residues was observed when the temperature decreased to 10 °C **(Figure S4B)**, which indicates a similar formation of cation-π structure with condensates prepared using the monomer unit (**Figure S4C**). Notably, with decreasing temperature, a shoulder peak emerged from the resonance of Tyr **(Figure S4B)**, which confirms that different conformational arrangements coexisted within the condensates. These heterogenous structures suggest that the multiple-step conformational changes occurred as responses to the temperature-dependent transition.

We next implemented fluorescence recovery after photobleaching technique (FRAP) to evaluate whether transition pathways could encode mesoscopic physical properties of condensates formed by full-length RLP_WT_ by doping a 50:1 molar ratio of RLP_WT_-mEGFP fusion protein as a fluorescent label. The final conditions of the condensates formed by distinct pathways are identical in terms of protein concentration, buffer condition and environmental temperature. Compared to condensates formed by temperature-dependent transition, condensates formed by decreasing ionic strength showed an apparent slower recovery process (**Figure 5B**). However, we did not find a noticeable difference in the fraction of recovery after equilibrium. This observation confirms that the pathway can indeed modulate the physical properties of condensates, suggesting that the feature that recognizes in a peptide monomer unit can reflect on the overall property of the full length homopolymer.

### Pathway defines the microenvironments of condensates and their electrochemical functions

Next, we focused on investigating whether the pathway could serve as the key factor modulating the microenvironments of condensates^18,25,68–70^, which have emerged as the key factor defining the functions of condensates. To this end, we first analyzed whether the interior pH of condensates could be defined by the transition process using RLP_WT_ condensates formed by distinct transition processes. Still, the final solution condition of the RLP_WT_ condensates was the same (see Methods for details). We employed the well-established ratiometric C-SNARF-4 assay^19^, in which the ratio of fluorescence generated from two distinct emission channels can be used to imply pH. We found that under the same bulk environmental pH, compared to condensates formed by decreasing temperature, condensates formed by decreasing ionic strength showed a more basic pH (**Figure 5D**). This might correlate with the nature of pi-based interactions, which are the key difference in the dense phase interactions between these two pathways. These pi-based interactions essentially define the asymmetry of availability of the charged residues or ions in the dense phase. This can possibly achieve by multiple scenarios: 1) cation-pi interaction leads to an available anion for attracting ions to realize neutral condition; 2) pi-pi interaction leads to polarized attraction of anion or cation to alter the electrostatic interaction or attract additional ion due to polarity^71^; 3) charge-dependent polarization effect induced the change of local water dissociation constant. These scenarios can all alter the dense phase apparent pH condition. These diverse possibilities further emphasize the importance to understand pathway-dependent sequence grammar and environmental effect on condensate properties. The differences of pH gradient suggest a difference in interphase Galvani potential^24,25^. This interphase potential difference can possibly manifest as a difference in interfacial electric field^19^. We used the ANEPPS assay^19,69^ to evaluate the change of interfacial field and observed a slight difference following the same trend as pH gradient of condensates prepared through distinct pathways (**Figure S5B)**. This observation confirms that pathway can encode distinct electrochemical environments of condensates.

As the interphase electric potential and the interfacial field are the underlying drivers of the inherent redox activity, as widely observed in the case of microdroplet chemistry^72–76^, we hypothesized that the pathway can further mediate distinct electrochemical functions of condensates^24,65^. To this end, we employed previously established fluorogenic assays to probe the capability of individual condensates on driving redox reactions^24,65^. Specifically, we implemented the resazurin assay to study the reduction reaction and the PY-1 assay to study the oxidation reaction^19,77^. Indeed, we found that the condensates formed through distinct pathways showed distinct electrochemical activity (**Figure 5E**). The capability to drive electrochemical reaction aligns with their corresponding electric potential gradient. This observation emphasizes that transition pathway serves as a critical element dictating the functions of biomolecular condensates.

## Discussion

The formation of biomolecular condensates can be driven by distinct phase transition pathways, as exemplified by temperature-dependent and osmosis-dependent condensate formations in living cells. A typical phase transition drivers can form condensates through these two distinct pathways, such as FUS, hnRNP A1 and DDX4 proteins^31,56,78^. However, whether cells can utilize pathway to encode different transition driving forces in order to regulate the properties and functions of condensates is unknown. We herein show that for the same IDP, temperature-dependent or osmosis-dependent phase transitions are driven by distinct molecular driving forces, which encode different dense phase interactions. A notable finding is that in the temperature-dependent transition pathway, phase transition is initiated by a specific interaction at a specific temperature, which is an anion-modulated cation-pi interaction, leading to coil-to-globule transition. The dense phase interactions further define the physical properties and the microenvironments of the condensates, which in turn encode distinct inherent electrochemical functions of condensates.

An IDP can undergo phase transition through distinct pathways. Our study shows that the environmental factors are coupled with distinct molecular interactions. This finding suggests that the general molecular grammars^15,28,31,32,79–81^ developed for understanding the phase behaviors of IDPs can further be expanded by considering the contributions of different amino acids under different environmental driving forces. From an experimental aspect, our finding implies that the procedures for preparing condensates can be critical to understand the properties and functions of condensates. As shown in a recent study^82^, the experimental order of adding components of a condensate can result in differences in condensate structure, which implies the importance of dynamics between different interactions. Our study further highlights that even for a single-component condensate, the homotypic interactions can be differentiated by the environmental driving forces of phase separation. Together, these findings highlight the contributions of distinct thermodynamic driving forces of phase transition on modulating the properties and functions of condensates.

Another interesting finding is that the UCST phase transition is initiated by the conformational change of one specific residue, Asp, within the RLP monomer. Asp exhibits a bistable state after crossing a specific temperature during transition, at which strong π-based interactions are established and lead to collapse transition of the peptide chain. Then, at a lower temperature, phase separation, as shown by changed diffusivity of peptide, happens. This process (**Figure 4E**) suggests that the temperature-induced transition is a multi-step process, through which a lower energy state is achieved. This observation suggests that understanding the temperature-dependent free energy hydration of each amino acids in a sequence-dependent manner^60,83^ can be important to define the key residues that are critical to initiate phase transition and define critical transition temperature. From a thermodynamic perspective of biomolecular phase transition, which typically exhibits enthalpy-entropy compensation behavior^61^, understanding the contributions of water reorganization enthalpy in a sequence- and temperature-dependent manners can further expand the sequence grammars of IDPs on encoding thermos-responsive phase behaviors^84^. This aspect can also be informative for IDP and condensate designs for synthetic biology applications^3^.

## Materials and methods

### Preparation of protein materials

GRGDSPYS monomer unit was synthesized and purchased from GenScript. For the preparation of full-length RLP_WT_ [GRGDSPYS]_20_ and RLP_WT_-mEGFP fusion, the plasmids were a gift from the Chilkoti lab from Duke University. We followed the detailed protocol as demonstrated in our previous studies for the expression and purification of this RLP_WT_^19,29^. After the typical inverse-temperature-dependent cycle for purification, the proteins were isolating with AKTA size exclusion chromatography (cytivia) and the final purity of the protein is higher than 95%. We stored the purified protein by flash frozen in liquid nitrogen and stored in -80 °C.

### Phase separation of RLP_WT_ through distinct pathways

The phase separation of the RLP_WT_ was reported as previous work. RLP_WT_ protein solution was premixed with 5% of RLP_WT_-mEGFP in the stock solution buffer. For the temperature-induced pathway, the 70 µL dilute buffer solution (50 mM Tris-HCl, pH=7.5) was mixed with the 30 µL protein solution (50 mM Tris-HCl, 500 mM NaCl, pH=7.5) to a final RLP_WT_ concentration of 20 µM at 37°C and incubated for 30 min, at which no phase separation was observed. The solution was then transferred to 25°C environments for 15 min. For the osmosis-dependent pathway, the 70 µL dilute buffer solution (50 mM Tris-HCl, pH=7.5) was mixed with the 30 µL protein solution (50 mM Tris-HCl, 500 mM NaCl pH=7.5) to a final RLP_WT_ concentration of 20 µM and incubated at 25°C for 45 min. The condensates were then subjected for characterizations.

### Preparation of peptide condensates

For temperature-dependent pathway, the solution containing 70 mg/ 50 µL of the peptide in 50 mM MES 0.1 mM NaCl (pH 5.9) was incubated at 37°C for 30 min and then incubated at 4°C. Then, the condensates were separated through centrifugation and the dense phase was immediately frozen in liquid nitrogen and lyophilized for solid-state NMR characterizations.

For osmosis-dependent pathway, the 10 µL dilute buffer solution (50 mM MES, pH=5.9) was added to the 90 µL peptide stock solution to realize a final peptide concentration of 70 mg/50 µL, 50 mM MES, 4 M NaCl (pH=5.9) at room temperature. After the solution becomes turbid, the dense phase was separated, and the sample was put in liquid nitrogen immediately to freeze the condensates and lyophilized for solid-state NMR characterizations.

### Morphology characterization

Cryo-transmission electron microscopy (cryo-TEM) images were acquired using a JEOL JEM-2310 instrument. Micrographs were recorded using a CCD camera from Gantega (Olympus Soft Imaging Solutions). Each sample was observed under the amorphous ice layer at 77K in case of damage from the electron beam. The Leica DMi8 was used for the fluorescence recovery after photobleaching.

### Spectroscopy characterization. ESI-MS

ESI-MS spectrum was obtained using a Bruker APEX II-FT-ICR-MS.

0.1 mg peptide was dissolved into the MilliQ water (1 mL) and the ESI spectrum was obtained in the positive mode and the scan number is 20. The full scan single mass spectra were obtained by scanning in a range of m/z = 58-1000. Calculated molecular weights refer to the m/z values given by the data analysis software.

### Solution-state NMR for RLP monomer

All experiments were carried out with 20 mg/mL peptide in 50 mM MES, pH 5.9 and 10% D_2_O for NMR field-frequency lock. All NMR experiments were recorded on the Bruker Avance 700 MHz spectrometer equipped with a triple resonance cryogenic probe. All chemical shifts from NMR were referenced to TMS (δH=0 ppm) and dichloromethane (δC=53.48 ppm) All experiments were processed and visualize using BRUKER Topspin version 4.1.3 software.

Pulsed field gradient NMR diffusion experiments on peptide with different temperatures (4°C, 10°C, 15°C, 20°C, 25°C, 30°C, 35°C, 37°C, and 40°C) were performed as pseudo-2D experiments using the standard Bruker ledbpgppr2s pulse sequence with a gradient strength from 3.2–59.9%. Data were analyzed by integrating resonances corresponding to regions including aspartic acid, arginine and tyrosine side chains in peptide GRGDSPYS (regions from 9.6–6.4 ppm) and the aliphatic region (4.4–1.5 ppm) for lysozyme. Data were analyzed for buffer signals (MES) from peptide for regions 3.379 and 3.584 ppm, respectively.

^1^H-^15^N Heteronuclear Single Quantum Coherence (HSQC) for peptide sample was acquired with 128 and 2048 total points in the indirect ^15^N and direct ^1^H dimensions by using pulse sequence hsqcf3pph19 at 37℃, with acquisition times of 143 and 24 ms and spectral widths of 37.0 ppm and 10.2 ppm, centered at 4.68 ppm and 116.99 ppm, respectively.

^1^H-^15^N Heteronuclear Multiple Bond Correlation (HMBC) for peptide sample was acquired with 128 and 2048 total points in the indirect ^15^N and direct ^1^H dimensions by using pulse sequence hmbcgpndqf at 37℃, with acquisition times of 27.1 and 155.6 ms and spectral widths of 58.2 ppm and 8.2 ppm, centered at 5.17 ppm and 105.65 ppm, respectively.

^1^H-^13^C Heteronuclear Single Quantum Coherence (HSQC) for peptide sample was acquired with 2048 and 1024 total points in the direct ^1^H and indirect ^13^C dimensions by using pulse sequence hsqcf3pph 19 at 37℃, with acquisition times of 163.8 and 17.1 ms and spectral widths of 8.9 ppm and 170.0 ppm, centered at 4.7 ppm and 98.51 ppm, respectively.

^1^H-^13^C Heteronuclear Multiple Bond Correlation (HMBC) for peptide sample was acquired with 4096 and 1024 total points in the direct ^1^H and indirect ^13^C dimensions by using pulse sequence hmbcgpl2ndwg at 37℃, with acquisition times of 327.7 and 17.1 ms and spectral widths of 8.9 ppm and 170.0 ppm, centered at 4.7 ppm and 98.51 ppm, respectively.

^1^H-^1^H Total Correlation Spectroscopy (TOCSY) for peptide sample was acquired with 8192 and 128 total points in the direct ^1^H and direct ^1^H dimensions by using pulse sequence dipsi2gphh19 at 37℃, with acquisition times of 499.7 and 7.8 ms and spectral widths of 11.71 ppm and 11.70 ppm, centered at 4.68 ppm and 4.68 ppm, respectively.

^1^H-^1^H Nuclear Overhauser Effect (NOE) for peptide samples were recorded with a mixing time of 500 ms with 1024 and 1024 total points in the direct ^1^H and direct ^1^H dimensions by using pulse sequence noesygpph19 at 37℃ and 4℃, respectively, with acquisition times of 499.7 and 7.6 ms and spectral widths of 11.71 ppm and 12.00 ppm, centered at 4.68 ppm and 4.68 ppm, respectively.

### Solid-state NMR for RLP monomer

The *in-situ* temperature-changing solid-state NMR spectra were recorded at 14.1 T (599.2 MHz ^1^H NMR frequency) static magnetic-field strength by using a Bruker AVANCE III 600. The spectrometer has a magic angle spinning (MAS) 4.0 mm rotor at an MAS rate of 12 kHz of 1D ^13^C cross-polarization measurements.

The 2D ^1^H-^13^C Heteronuclear Correlation (HECTOR) spectra for peptide samples were recorded with different contact time (0.1 ms, 0.5 ms, 1 ms, and 2 ms) with 2048 and 64 total points in the indirect ^13^C and direct ^1^H dimensions by using pulse sequence lghetfq at 25℃, with acquisition times of 22.5 and 0.7 ms and spectral widths of 301.5 ppm and 70.7 ppm, centered at 99.18 ppm and 2.85 ppm, respectively.

All chemical shifts from ssNMR were referenced to adamantane (δH=1.91 ppm, δC=38.48 ppm). Compressed air was used to regulate the sample’s temperature at room temperature. All sample temperatures were checked by measuring the chemical shift of ^79^Br in KBr powder. A linear fit of the ^79^Br data yields a −0.0249 + 0.0015*ppm*/(*T* − 273.15)

### Solution-state NMR for full-length protein RLP_WT_

The unlabeled RLP_WT_ protein at the concentration of 75 µM was carried out at different temperature (37℃, 30℃, 20℃, 10℃ and 4℃) in 150 mM PB buffer, pH=7.5 and 10% D2O for NMR field-frequency lock. And all NMR experiments were performed on 600 MHz spectrometer Bruker Avance III 600MHz with a quadra resonance cryogenic probe.

*In-situ* temperature-changing ^1^H spectra and ^1^H-^1^H DOSY experiments were conducted under the PB buffer at pH 7.5 (150 mM phosphate buffer). The 1D temperature-changing ^1^H experiment was employed at the 64 scans at the target temperature with the pulse sequence p3919pg.

Pulsed field gradient NMR diffusion experiments on peptide with different temperatures (37°C, 30°C, 20°C, 10°C and 4°C) were performed as pseudo-2D experiments using the standard Bruker stebpgp1s19 pulse sequence with a gradient strength from 3.2–99.9%. Data were analyzed by integrating resonances corresponding to regions including aspartic acid, arginine and tyrosine side chains in WT20 (regions from 8.4–5.2 ppm). NMR spectra were recorded after the sample was maintained at the desired temperature for 30 minutes, ensuring thorough data collection. The data were then meticulously processed using BRUKER Topspin version 4.1.3.

### Pathway-dependent condensate formation for RLP_WT_

For the temperature-induced pathway, stock solutions (RLP_WT_ in 50 mM Tris, 500 mM NaCl, pH 7.5) and dilute solutions (50 mM Tris, 0 mM NaCl, pH 7.5) were incubated at 42°C for 30 minutes. Then, 350 µL of dilute buffer was mixed with 150 µL of protein solution to achieve a final RLP_WT_ concentration of 20 µM at 42°C, followed by an additional 30-minute incubation. The samples were then transferred to room temperature (25°C) to induce temperature-dependent phase separation.

For the osmosis-induced pathway, the same stock and dilute solutions were incubated at 25°C for 30 minutes. Then, 350 µL of dilute buffer was mixed with 150 µL of protein solution, maintaining a final RLP_WT_ concentration of 20 µM at 25°C. This initiated osmosis-dependent phase separation.

To minimize batch-to-batch variation, RLP_WT_ was taken from the same stock solution for both pathways.

### Fluorescence recovery after photobleaching analysis of RLP_WT_ condensates formed through distinct pathways

The dynamics of RLP_WT_ condensates were characterized through FRAP using LECIA Stellaris 8 FELCON. The characterizations were conducted in an environmental chamber set at 25 ℃. The excitation was set at 488 nm and the Hyd detector was set at 500-540 nm to viasualize the condensates. For bleaching experiments, the laser power was set to 100% and the selected ROIs were bleached for 5 times before tracking the recovery of the signal under a laser intensity of 5 % and the detector gain set at 100%. The FRAP recovery was analyzed with LEICA LAS X life science software.

### C-SNARF-4 assay to evaluate the apparent pH value of condensates

C-SNARF-4 assay was used to assess the pH of condensates formed through different pathways. C-SNARF-4 dye was added to the samples at a final concentration of 2 μM with a 1:100 volume ratio. Samples were incubated in PhenoPlate™ 384-well microplates (Revvity) for confocal microscopy, and the dye was introduced 30 minutes before imaging to allow equilibration and accurate pH representation within condensates. Fluorescence signals were acquired using confocal microscopy with an excitation wavelength of 490 nm and emission detected at 540–590 nm (channel 1) and 610–660 nm (channel 2). ImageJ was used for fluorescence signal analysis, with regions of interest (ROIs) selected to encompass entire condensates. The fluorescence intensity ratio, calculated as the mean fluorescence per pixel in channel 1 divided by that in channel 2, was used to determine pH.

### Di-4-ANEPPS assay to evaluate the interfacial electric field of condensates

Di-4-ANEPPS assay was used to evaluate the interfacial electric field in condensates formed via different pathways. Di-4-ANEPPS dye was added to the samples to a final concentration of 1 μM. After incubation, samples were transferred into PhenoPlate™ 384-well microplates (Revvity) for confocal microscopy. The dye was introduced 10 minutes prior to imaging to ensure adequate incorporation. Fluorescence signals were acquired using confocal microscopy with an excitation wavelength of 470 nm and emission collected at 535–545 nm (channel 1) and 610–640 nm (channel 2). ImageJ was used to quantify fluorescence intensities, with regions of interest (ROIs) selected to encompass entire condensates. The fluorescence intensity ratio between the two channels, calculated as the mean fluorescence per pixel in channel 1 divided by that in channel 2, was used to determine interfacial electric field.

### PY1 assay for the characterization of H_2_O_2_ produced by condensates

Fluorogenic PY1 assay was used to measure the H₂O₂ production rate in condensates formed through different pathways. PY1 was added to samples at a final concentration of 20 μM and incubated for 30 minutes before imaging. Samples were then transferred into PhenoPlate™ 384-well microplates (Revvity) with a 40 μL sample volume per well for confocal microscopy. Fluorescence signals were acquired with an excitation wavelength of 514 nm, and emission was detected at 538–558 nm. ImageJ was used for fluorescence signal analysis, with regions of interest (ROIs) selected to encompass entire condensates.

### Resazurin assay for the characterization of electron donating capability by temperature-induced and osmosis-induced condensates

Resazurin was used to assess the reducing capacity of condensates formed through different pathways. Resazurin was added to samples at a final concentration of 1 mM and incubated for 30 minutes in PhenoPlate™ 384-well microplates (Revvity) before imaging. Fluorescence signals were acquired using confocal microscopy with an excitation wavelength of 532 nm, and emission was detected at 540–590 nm. ImageJ was used for fluorescence signal analysis, with regions of interest (ROIs) selected to encompass entire condensates.

## Acknowledgements and funding sources

We acknowledge the experimental and funding supports from the Center for Biomolecular Condensates at Washington University in St. Louis.

## Author Contributions

Y.D. devised the idea. X.R. and Y.D. designed the experiments. X.R., L.Y., M.W.C. performed and analyzed the experiments. Y.D. and X.R. wrote the manuscript.

## Notes

### Competing Interest Statement

The authors have declared no competing interest.

