## Supplementary File for "Phase transition pathways encode distinct physicochemical properties of biomolecular condensates"

**This PDF file includes:**
**Figure S1 to S5**

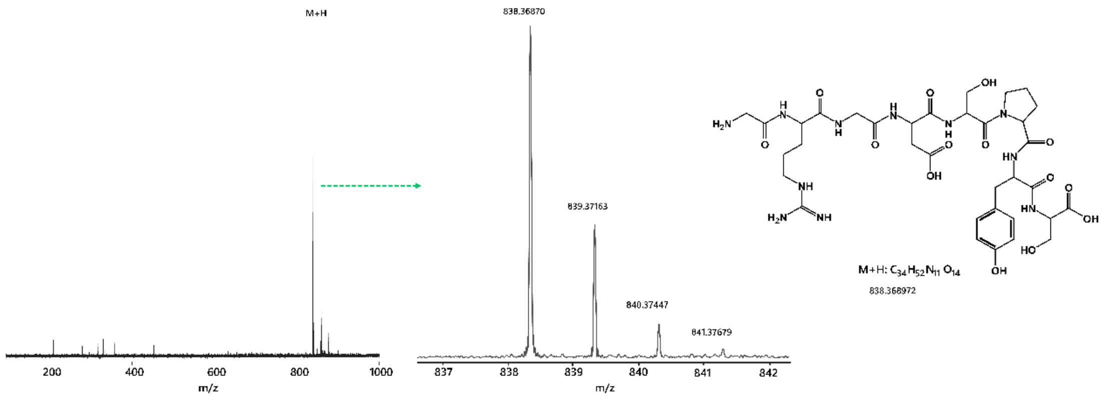


**Figure S1. HR-ESI mass spectroscopy analysis of the peptide unit.**

The HR-ESI mass spectra confirms that the synthesis of the repeat unit peptide GRGDSPYS.

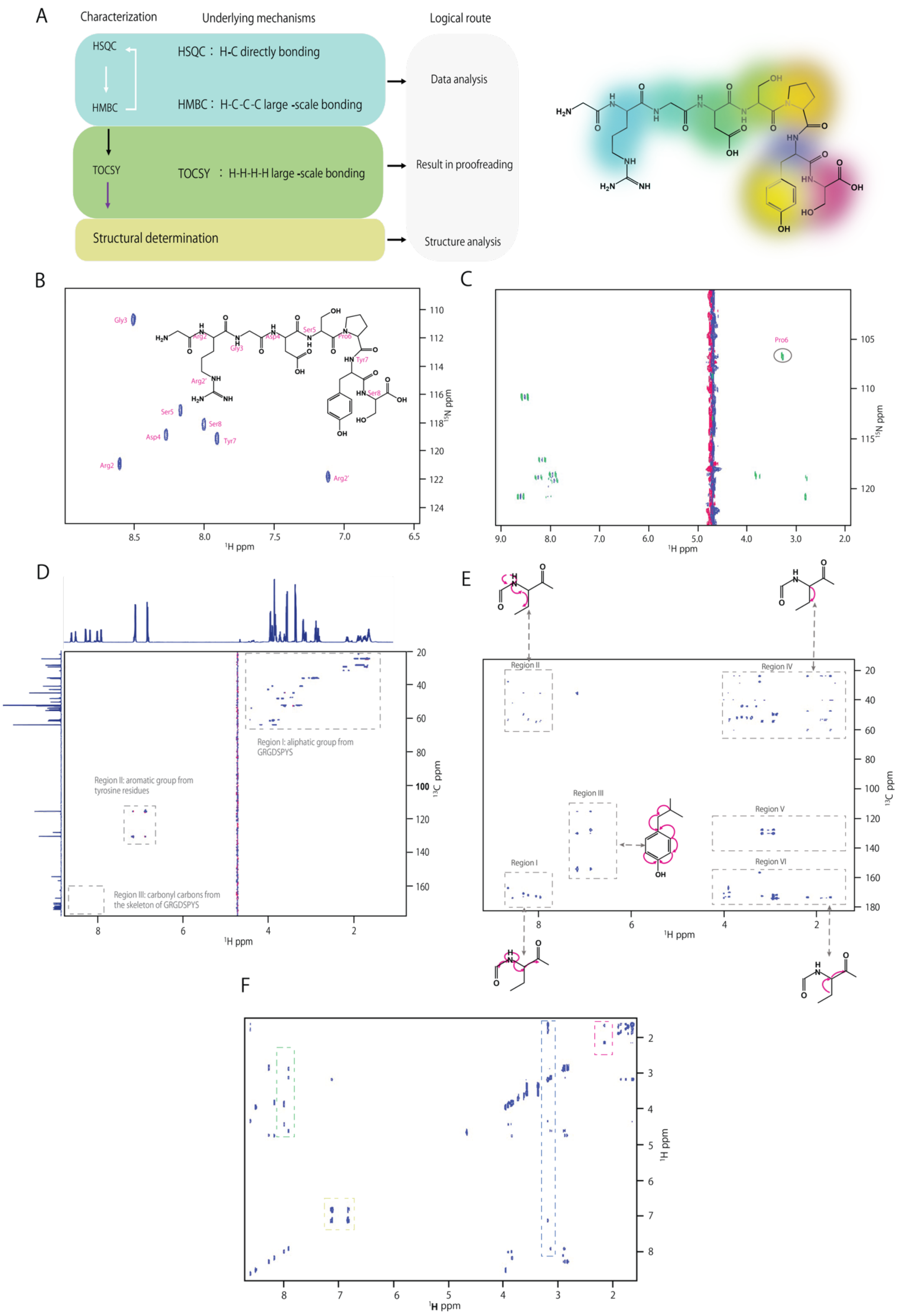
A
Characterization
Underlying mechanisms
Logical route
HSQC
HMBC
HSQC： H-C directly bonding
HMBC： H-C-C-C large -scale bonding
Data analysis
TOCSY
TOCSY ： H-H-H-H large -scale bonding
Result in proofreading
Structural determination
Structure analysis
B
Gly3
Ser5
Ser8
Asp4
Tyr7
Arg2
Arg2'
¹H ppm
¹⁵N ppm
C
Pro6
¹H ppm
¹⁵N ppm
D
Region I: aliphatic group from GRGDSPYS
Region II: aromatic group from tyrosine residues
Region III: carbonyl carbons from the skeleton of GRGDSPYS
¹H ppm
¹³C ppm
E
Region II
Region IV
Region III
Region V
Region I
Region VI
¹H ppm
¹³C ppm
F
¹H ppm
¹H ppm

**Figure S2. NMR characterizations of the structural information of the monomer RLP unit.**

**A,** NMR characterizations and the logical route for the construction of atomic level chemical shifts for the monomer RLP unit.

**B,** 2D $^{1}H$-$^{15}N$ HSQC was used to determine the attribution of NMR signals on hydrogen-linked nitrogen atoms.

**C,** 2D $^{1}H$-$^{15}N$ HMBC was used to determine the attribution of NMR signal on the nitrogen atom of the Pro residue.

**D,** 2D $^{1}H$-$^{13}C$ HSQC was used to determine the attribution of NMR signals of carbon atoms on the peptide side chain.

**E,** 2D $^{1}H$-$^{13}C$ HMBC was used to determine the attribution of NMR signals of carbon and hydrogen atoms on the whole peptide.

**F,** 2D $^{1}H$-$^{1}H$ TOCSY was used to check the chemical shift attribution of the whole molecular structure of peptide.

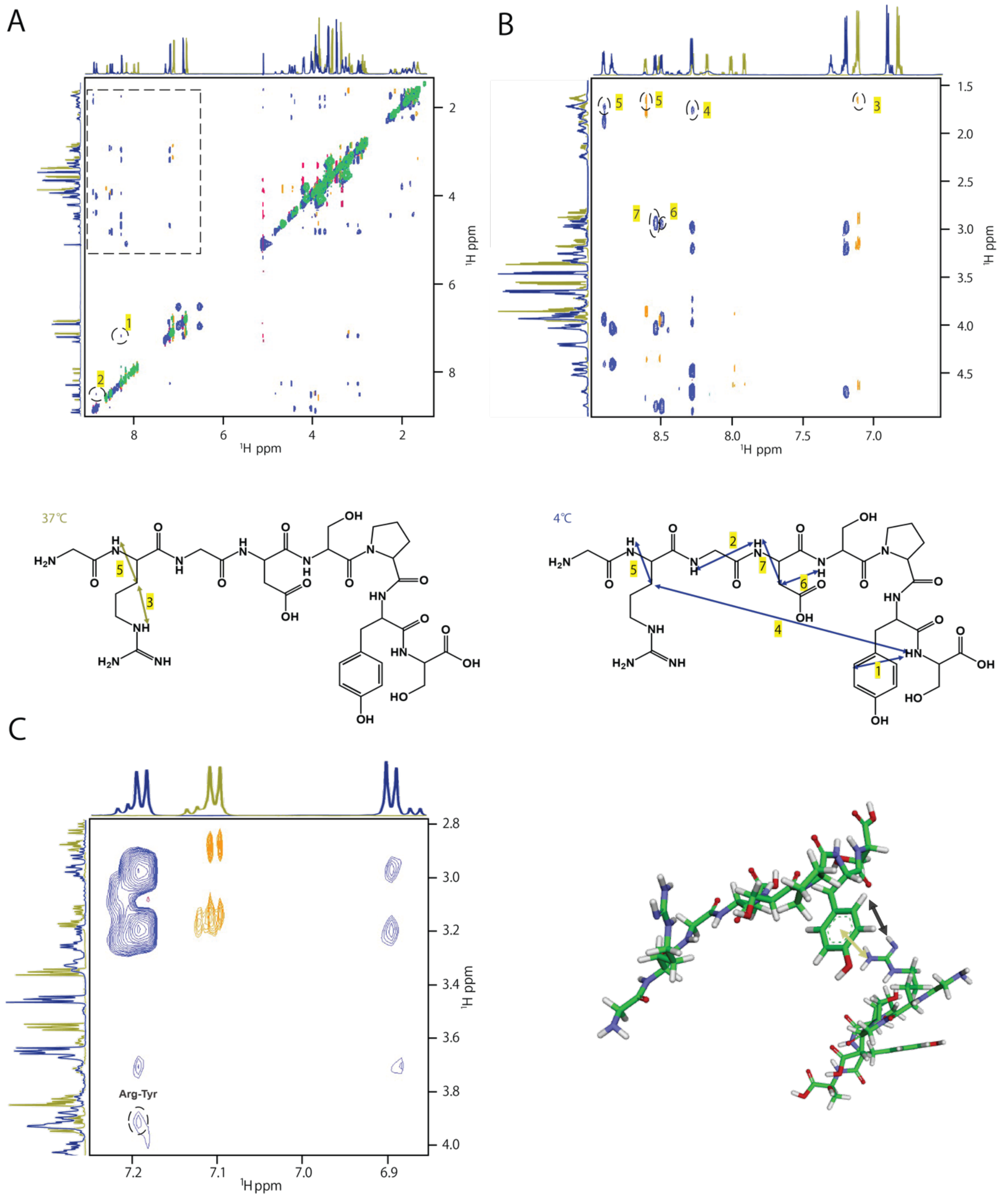


**Figure S3. 2D $^1$H-$^1$H NOE spectra of GRGDSPYS at different temperatures (4°C and 37°C).**

**A,** NOE spectra exhibited the two geometric distances between carbon atoms to their responding hydrogen atoms (marked as 1 and 2) detected at 4°C.

**B,** Selected regionS of the NOE spectra exhibited the four different distances (marked as 4,5,6 and 7) that can be detected at 4°C and two different distances (marked as 3 and 5) at 37°C. Sample data acquired at 37°C was exhibited by yellow (positive signal) and green (negative signal) cross peak dots, while the data acquired at 4°C was exhibited by blue (positive signal) and red (negative signal). The schematic illustrations of these different distances (1, 2, 3, 4, 5, 6 and 7) were exhibited below the NOE spectra.

**C,** 2D $^{1}H$-$^{1}H$ NOE indicated the cation-π interactions between Arg (R) and Tyr (Y) formed within the molecule at 4°C.

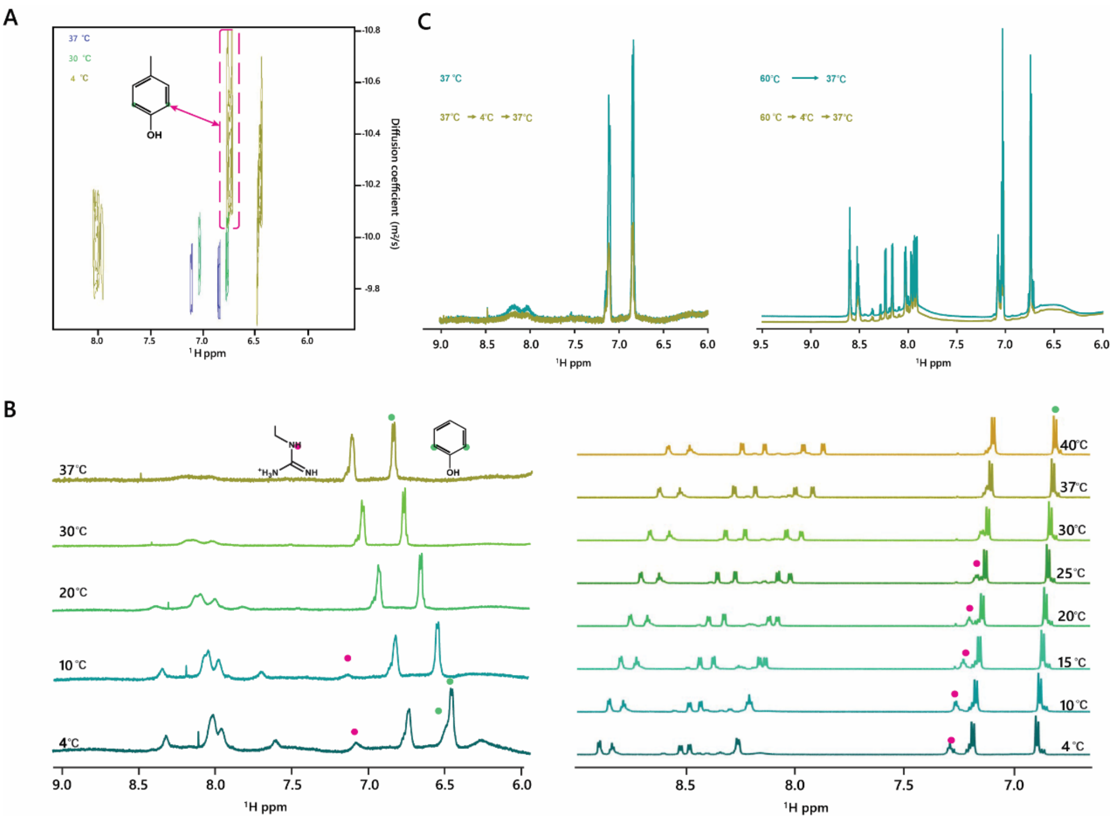


**Figure S4. $^{1}H$ NMR characterization of full-length RLP$_{WT}$.**

**A,** *In-situ* temperature-changing diffusion-ordered spectroscopy (DOSY) analyzes the proton of RLP$_{WT}$ upon decreasing temperature. The ortho proton atoms of Tyr residues exhibited a broad diffusion coefficient, which suggests that tyrosine residues organized heterogenous assemblies with different sizes within condensates.

**B,** 1D temperature-changing $^{1}H$ NMR spectra shows that the phase transition is driven by cation-π interaction between Arg (R) and Tyr (Y). These features are identical between the full-length protein RLP$_{WT}$ and monomer unit (right).

**C,** *In-situ* temperature-changing 1D $^{1}H$ spectra of full-length protein RLP$_{WT}$ and monomer unit (right). A similar trend in the intensity profile was observed between the protein and the monomer unit peptide. This suggests that the protein maintains a partially tight packing structure from 4°C to 37°C. A similar trend is observed in the monomer unit peptide, confirming that interactions between Asp, Arg and Tyr are key factors in forming and structuring condensates formed by RLP$_{WT}$.

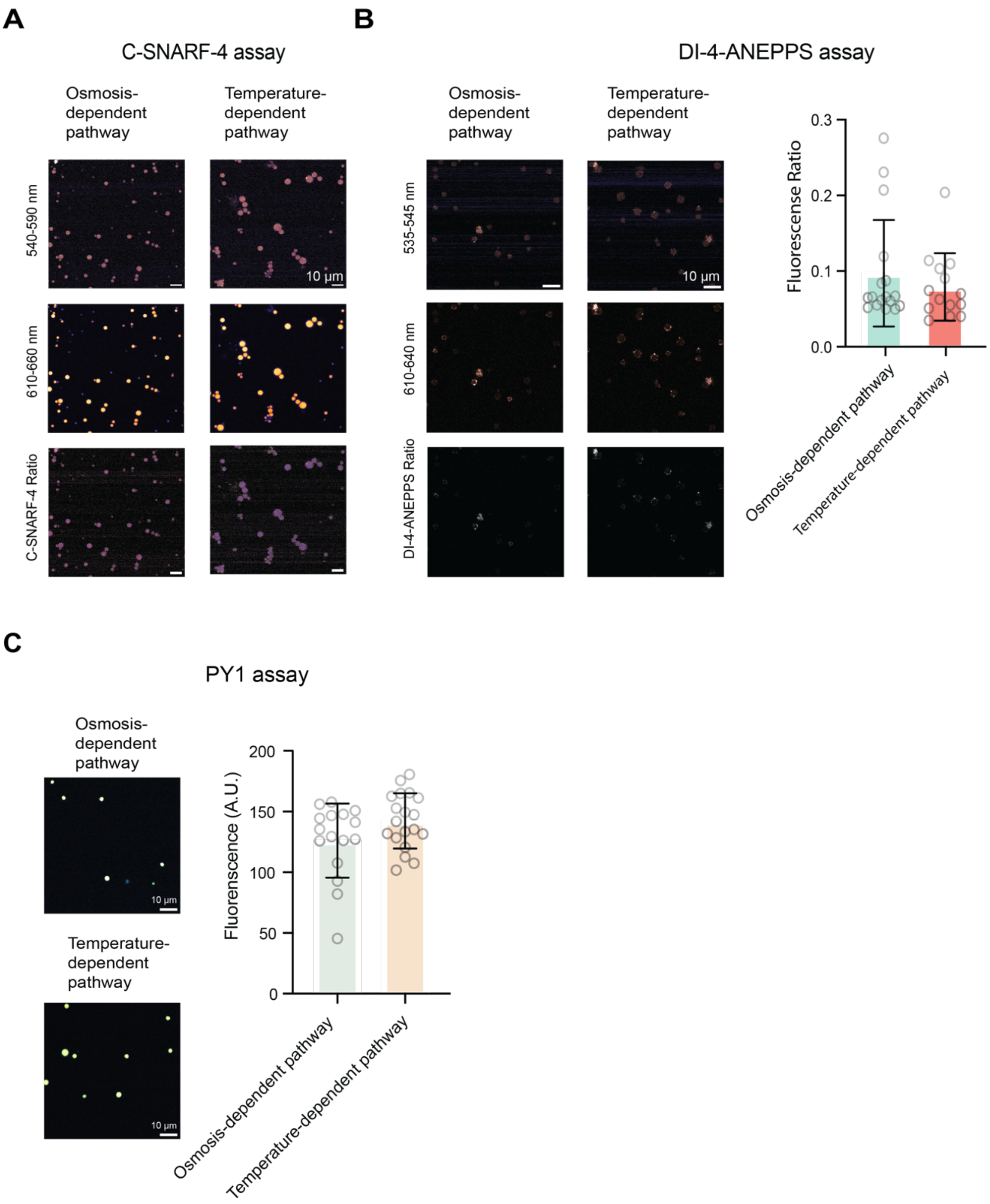


**Figure S5. Evaluation of pathway-dependent electrochemical environments and oxidation activity of biomolecular condensates formed by full-length $RLP_{WT}$.**

**A,** Representative images of C-SNARF-4 assay measurement of apparent pH in the dense phase of condensates formed through different pathways. Scale bar: 10 µm

**B,** Representative images of Di-4-ANEPPS assay for the evaluation of the interfacial potential of condensates formed through different transition processes. Each data point represents an individual condensate. Scale bar: 10 µm

**C,** PY1 assay analysis of spontaneous oxidation activity of condensates based on $H_2O_2$ production. Each data point represents an individual condensate. Scale bar: 10 µm
